# Individual differences in oxycodone addiction-like behaviors in a large cohort of heterogeneous stock (HS) rats

**DOI:** 10.1101/2022.07.26.501618

**Authors:** Marsida Kallupi, Giordano de Guglielmo, Lieselot LG Carrette, Sierra Simpson, Jenni Kononoff, Adam Kimbrough, Lauren C Smith, Kokila Shankar, Alicia Avelar, Dana Conlisk, Molly Brennan, Lani Tieu, Sharona Sedighim, Brent Boomhower, Lisa Maturin, McKenzie J Fannon, Angelica Martinez, Caitlin Crook, Selen Dirik, Nathan Velarde, Paul Schweitzer, Selene Bonnet-Zahedi, Dyar N. Othman, Benjamin Sichel, Kwynn Guess, Beverly Peng, Andrew S. Hu, Lucas E. Chun, Kristel Milan, Justin Lau, Yicen Zheng, Ashley Vang, Leah C. Solberg Woods, Abraham A. Palmer, Olivier George

## Abstract

Family and twin studies demonstrate that genetic factors determine 20-60% of the vulnerability to opioid use disorder. However, the genes/alleles that mediate the risk of developing addiction-related behaviors, including the sensitivity to the analgesic efficacy of opioids, the development of tolerance, dependence, and escalation of oxycodone taking and seeking, have been ill-defined, thus hindering efforts to design pharmacological interventions to enable precision medicine strategies. Here we characterized oxycodone addiction-like behaviors in heterogeneous stock (HS) rats, that show high genetic diversity that mimics the high genetic variability in humans. HS rats were allowed to self-administer oxycodone for two h/daily for four days (ShA) and then moved to 12h/daily (LgA) for 14 days. Animals were screened for motivation to self-administer oxycodone using a progressive-ratio (PR) schedule of reinforcement and for the development of withdrawal-induced hyperalgesia and tolerance to the analgesic effects of oxycodone using the von-Frey and tail immersion tests, respectively. To reduce cohort-specific effects, we used cohorts of 46-60 rats and normalized the response level within cohorts using a Z-score. To take advantage of the four opioid-related behaviors and further identify subjects that are consistently vulnerable *vs*. resilient to compulsive oxycodone use, we computed an Addiction Index by averaging normalized responding (Z-scores) for the four behavioral tests. Results showed high individual variability between vulnerable and resilient rats, likely to facilitate the detection of gene variants associated with vulnerable *vs*. resilient individuals. Such data will have considerable translational value for designing follow-up studies in humans.

## Introduction

More than 2 million Americans currently suffer from substance use disorders related to prescription opioid pain relievers, including oxycodone (Oxycontin^®^, Roxycodone^®^, Oxecta^®^), and 500,000 are addicted to heroin. (1). Over the past 15 years, the consumption of oxycodone increased by ~500%, and opioid-related overdose deaths quadrupled (2, 3). Although opioid medications effectively treat acute pain and help relieve chronic pain for some patients (4), the risk of addiction presents a dilemma for healthcare providers who seek to relieve suffering while preventing drug abuse and addiction. Little is known about the risk for addiction among those treated for chronic pain and how basic pain mechanisms interact with prescription opioids to influence addiction potential. The vulnerability to substance abuse and addiction, including oxycodone use disorder, have a heritable component. Family and twin studies demonstrate that 20-60% of the vulnerability to opioid use disorder is determined by genetic factors (5–7). Many genetic factors interacting together can contribute to the vulnerability or the resilience to the opioid use disorder, making the genetic contribution polygenic. In recent years, human genomewide association studies (GWAS) have expanded dramatically in size and sophistication, which has led to numerous loci being implicated in a range of addiction-related traits (8–12); however, most of these loci do not replicate between studies and the vast majority of genes underlying the vulnerability to opioid use disorder remain unknown. Clinical studies on oxycodone use disorder face significant problems given the inaccuracy of self-reported addiction-related measures, the lack of extensive analyses of behavior, the low levels of biological samples and the missing control of possible environmental factors contributing to opioid use disorder. Although qualitative reports are highly relevant for diagnosing oxycodone use disorder (DSM-5), they are the least preferred approach for GWAS because they lead to low power for detecting genetic effects (13).

Therefore, the genetic underlying individual differences of oxycodone addiction-like behaviors are still poorly understood. (14, 15). Together with the progress in human genetics, the methodology for GWAS in mice and rats has also improved over the past decade. Previous studies were characterized by strains with limited recombination, inadequate genotyping platforms, and poor statistical approaches. An important step was the use of animal populations in which inbred founders were characterized by several generations of recombination, allowing the fine mapping of associated loci. The improvement of the genotyping techniques and the statistical approaches allowed to perform GWAS in laboratory animals to identify small genetic intervals (one or a few genes).

A significant achievement in the field has been the recent development of the extended access to oxycodone self-administration model that provided an animal model with high face, predictive, and construct validity for opioid use disorder (16, 17). Another reason for the failure of drug development for oxycodone use disorder is the lack of knowledge of gene variants that are associated with compulsive oxycodone use. Candidate genes that are associated with heroin use disorder and analgesic effects of analgesic opioids (18) have been identified, including genes that are implicated in the pharmacodynamics (OPRM1, COMT) and pharmacokinetics (CYP2D6, CYP3A4/5, ABCB1) of opioids (18). However, such studies cannot consistently be replicated (14) and have focused on pharmacological targets that have already been well explored and thus do not provide novel therapeutics. Unfortunately, genome-wide association studies (GWAS) have not been performed for oxycodone use disorder, and none of the 112 single-nucleotide polymorphisms (SNPs) in 25 candidate genes (including OPRM1 and COMT) showed significant associations with opioid dose (including oxycodone) in the treatment of pain (19). Thus, human GWAS have not yet made significant advances in our understanding of oxycodone use disorder or the analgesic effects of oxycodone. Although qualitative reports are highly relevant to the diagnosis of oxycodone use disorder (DSM-5), it is the least preferred approach for GWAS because it leads to low power for detecting genetic effects (13).

Here we comprehensively characterized oxycodone addiction-like behaviors in over 530 heterogeneous stock (HS) rats. HS rats are a highly recombinant rat intercross and represent a powerful tool for genetic studies (20, 21).

These rats (sometimes referred to as N/NIH) were created in 1984 by intercrossing eight inbred rat strains for more than 95 generations. HS rats are highly recombinant, allowing the fine mapping of genetic loci to exact intervals, significantly decreasing the number of potential underlying causative genes within each region. (21). Such genetic diversity makes HS rats an ideal resource for performing studies that are analogous to human GWAS. The rats underwent extended access of oxycodone self-administration that was combined with behavioral characterization of escalation of intake, analgesic effect of oxycodone, motivation despite an exponential increase in the cost to obtain oxycodone, withdrawal-induced hyperalgesia, and tolerance to the analgesic effects of oxycodone. The results of the different behavioral tests were normalized and computed into an Addiction Index (22) that allowed to identify animals vulnerable or resilient to compulsive oxycodone use.

## METHODS

Detailed procedures can be found in the George lab protocol repository on protocols.io (https://www.protocols.io/workspaces/george-lab).

### Animals

HS rats (Rat Genome Database NMcwiWFsm #13673907) created to encompass high genetic diversity by outbreeding eight inbred rat strains (ACI/N, BN/SsN, BUF/N, F344/N, M520/N, MR/N, WKY/N and WN/N) were provided by Dr. Leah Solberg Woods (Medical College of Wisconsin, now at Wake Forest University School of Medicine; *n* = 539). Rats were shipped at 3-4 weeks of age, kept in quarantine for 2 weeks and then housed two per cage on a 12 h/12 h reversed light/dark cycle in a temperature (20-22°C) and humidity (45-55%) controlled vivarium with ad libitum access to tap water and food pellets (PJ Noyes Company, Lancaster, NH, USA). All the procedures were conducted in strict adherence to the National Institutes of Health Guide for the Care and Use of Laboratory Animals and were approved by the Institutional Animal Care and Use Committees of The Scripps Research Institute and UC San Diego.

### Drugs

Oxycodone (National Institute on Drug Abuse, Bethesda, MD) was dissolved in 0.9% sterile saline and administered intravenously at a dose of 150 ug/kg/infusion.

This dose of oxycodone was selected based on previous studies (22-24) and because it produces significant plasma oxycodone concentrations (40 ng/ml; (25).

### Intravenous Catheterization

Rats were anesthetized with vaporized isoflurane (1-5%). Intravenous catheters were aseptically inserted into the right jugular vein using the procedure described previously (22). Catheters consisted of Micro-Renathane tubing (18 cm, 0.023-inch inner diameter, 0.037-inch outer diameter; Braintree Scientific, Braintree, MA, USA) attached to a 90 degree angle bend guide cannula (Plastics One, Roanoke, VA, USA), embedded in dental acrylic, and anchored with mesh (1 mm thick, 2 cm diameter). Tubing was inserted into the vein following a needle puncture (22G) and secured with a suture. The guide cannula was punctured through a small incision on the back. The outside part of the cannula was closed off with a plastic seal and metal cover cap, which allowed for sterility and protection of the catheter base. Flunixin (2.5 mg/kg, s.c.) was administered as analgesic, and Cefazolin (330 mg/kg, i.m.) as antibiotic. Rats were allowed seven days for recovery prior to any self-administration. They were monitored and flushed daily with heparinized saline (10 U/ml of heparin sodium; American Pharmaceutical Partners, Schaumberg, IL, USA) in 0.9% bacteriostatic sodium chloride (Hospira, Lake Forest, IL, USA) that contained 52.4 mg/0.2 ml of Cefazolin.

### Behavioral Testing

#### Operant self-administration

Self-administration (SA) was performed in operant conditioning chambers (29 cm × 24 cm × 19.5 cm; Med Associates, St. Albans, VT, USA) that were enclosed in lit, sound-attenuating, ventilated environmental cubicles. The front door and back wall of the chambers were constructed of transparent plastic, and the other walls were opaque metal. Each chamber was equipped with two retractable levers that were located on the front panel. Each session was initiated by the extension of two retractable levers into the chamber. Drugs (oxycodone, 150 ug/kg/infusion in saline) were delivered through plastic catheter tubing that was connected to an infusion pump. The infusion pump was activated by responses on the right (active) lever that were reinforced on fixed ratio (FR) 1 schedule, with the delivery of 0.1 ml of the drug per lever press over 6 s followed by a 20-s timeout period that was signaled by the illumination of a cue light above the active lever and during which active lever presses did not result in additional infusions. Responses on the left inactive lever were recorded but had no scheduled consequences. Fluid delivery and behavioral data recording was controlled by a computer with the MED-PC IV software installed. Initially, rats were trained to self-administer oxydone in 4 short access (ShA) sessions (2h/day). At the end of the ShA phase the animals were exposed to 14 extended access sessions (LgA, 12 h/day, 5 days/week Monday–Friday) to measure escalation of drug intake. Self-administration sessions started at the beginning of the dark phase of the light/dark cycle. The animals had access to food during the 12-h sessions. A fresh block of wood (3×3×3cm) was placed daily in the self-administration boxes.

#### Progressive Ratio Testing

Rats were tested on a progressive ratio (PR) schedule of reinforcement at the end of each phase (once after ShA and once after LgA, see Fig 1 for detailed timeline), in which the response requirements for receiving a single reinforcement increased according to the following equation: [5e^(injection number x 0.2^)] – 5. This resulted in the following progression of response requirements: 1, 1, 2, 2, 3, 3, 4, 4, 6, 6, 8, 8, 10, 11, 12, 13, 14, 15, 16, 17, etc +1 till 50, 60, 70, 80. 90, 100. The breakpoint was defined as the last ratio attained by the rat prior to a 60-min period during which a ratio was not completed, which ended the experiment.

**Figure 1.**
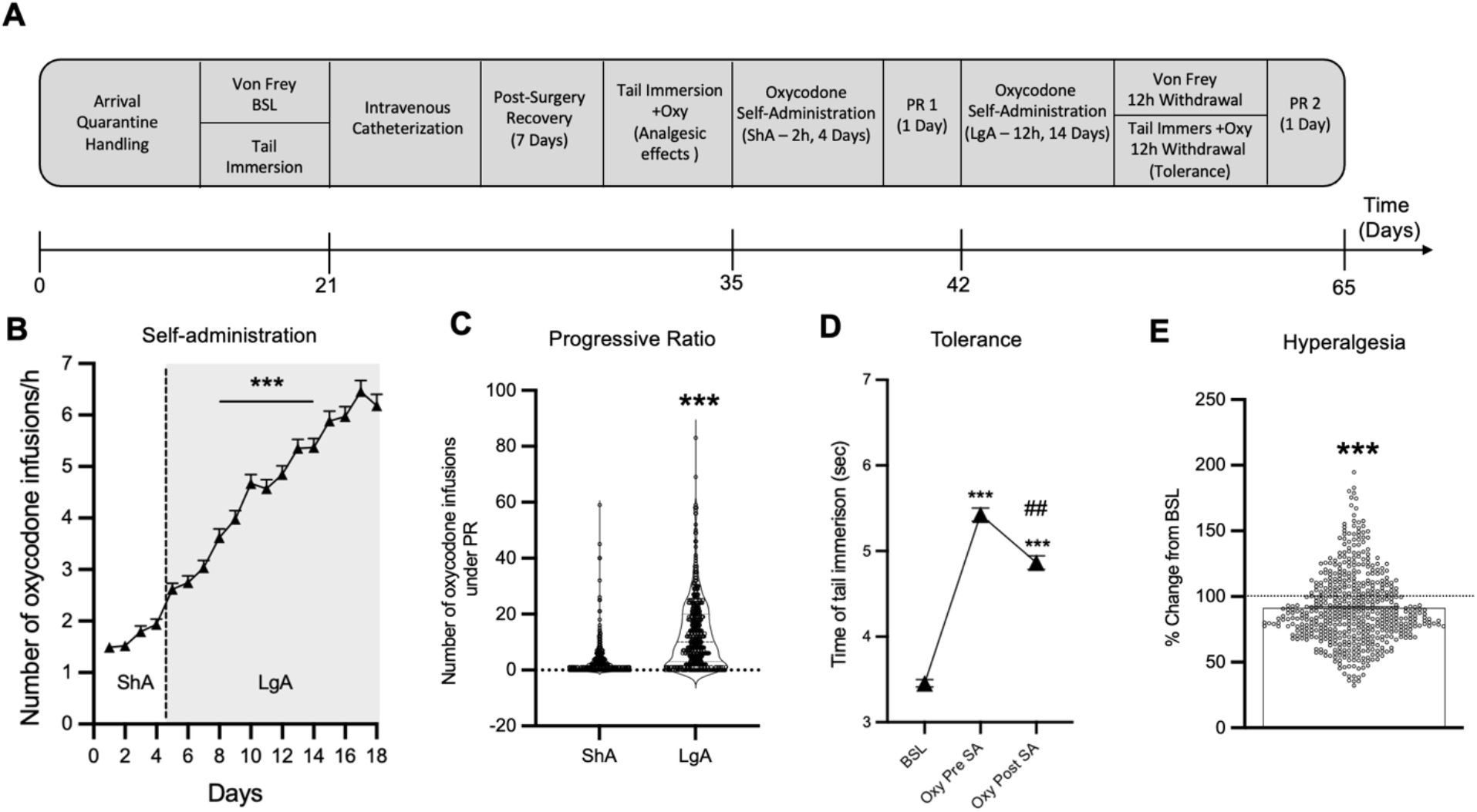
Individual differences in addiction-behaviors in oxycodone dependent HS rats. A) Timeline of the behavioral paradigms. B) Number of oxycodone infusions in the first hour of oxycodone self-administration during short (2h, ShA) and long (12h, LgA) access (*** p < 0.001 vs the first LgA session). C) Violin plot of number of oxycodone infusions under progressive ratio (PR) test after ShA and LgA (N= 523, *** p < 0.001). D) Development of tolerance to the analgesic effects of oxycodone. Tail withdrawal threshold at baseline, after 3 infusions of oxycodone pre- and post-extended access of oxycodone SA (*** p < 0.001 vs BSL and ### p < 0.001 vs Oxy pre-SA). E) Development of withdrawal induced hyperalgesia. Data are expressed as % change from baseline force in the Von Frey test. *** p < 0.001 vs BSL.

#### Mechanical Nociceptive Von-Frey Testing

Paw withdrawal in response to a dynamic plantar aesthesiometer (Ugo Basile) was tested for mechanical sensitivity as adapted from previous reports (26–28). Time of paw retraction and force exercised to produce a paw retraction was recorded for three measures for each back paw. Measurements were obtained at baseline (before any drug exposure), and during acute withdrawal (10-12h after the last oxycodone session) see Figs. 1 for timeline. Data are expressed as force (g) for baseline, and as percent change of force (g) from each individual baseline, to allow for a within-individual evaluation of hyperalgesia.

#### Analgesia by Tail immersion Testing

Rats were restrained in a soft tissue pocket, and the distal half of the tail was dipped into a water bath set at 52°C. The latency to withdrawal the tail from the water bath was measured. Two tail-withdrawal measures (separated by 30 s) were recorded and averaged. A 10 s cut-off time was applied to avoid tail burns (29). The tail immersion test was performed at three time points i) baseline levels (before any drug exposure), ii) before the initiation of the oxycodone ShA session. Rats received 2 infusions of oxycodone solution (150 ug/ml/kg) intravenously, and tail immersion test for measuring analgesia was performed 15 min after oxycodone administration, iii) after 14 LgA oxycodone sessions, 12 h into withdrawal from oxycodone. This measure was aimed to monitor the development of tolerance. Rats received 2 infusions of oxycodone solution (150 ug/ml/kg) 15 mins prior to the test.

#### Statistical Analyses

Data were analyzed using Prism 9.0 software (GraphPad, San Diego, CA, USA). Self-administration data were analyzed using repeated-measures analysis of variance (ANOVA) followed by Bonferroni post-hoc tests when appropriate. For pairwise comparisons, data were analyzed using the Student’s *t*-test. Correlations were calculated using Pearson *r* analysis. The data are expressed as mean ± SEM unless otherwise specified. Values of p < 0.05 were considered statistically significant.

## RESULTS

### Evaluation of Oxycodone Addiction-Like Behaviors in HS rats

We assessed addiction-like behaviors in HS rats self-administering oxycodone according to the standardized protocol shown in Figure 1A. Animals were tested in 14 cohorts with N=46-60 each. After catheterization surgery and recovery, the rats were trained to self-administer oxycodone in 4 daily 2h short access (ShA) sessions and then in 14 daily 12h long access (LgA) sessions. Over the course of the extended access of oxycodone self-administration phase, the animals showed escalation of intake, as measured by the significant increase of oxycodone rewards from day 4 of LgA onwards vs the first day of LgA (Fig. 1B, F_(17,8687)_ = 205.2 p < 0.0001 after one way ANOVA, followed by Bonferroni post hoc comparisons). Motivation for oxycodone was tested both after ShA and LgA, using a progressive ratio (PR) test (Fig. 1C). The number of oxycodone infusions significantly increased after extended access of oxycodone self-administration, compared to the levels at the end of the short access phase (t_538_ = 18.25, p < 0.001). The rats were also tested for their baseline response to pain, for their response to the analgesic effects of oxycodone and for the development of tolerance to the analgesic effects of oxycodone using the tail immersion test in 3 different phases of the behavioral protocol (See Fig. 1 A for detailed timeline of testing). As shown in the Figs. 1D, HS rats developed tolerance to the analgesic effects of oxycodone following extended access to oxycodone self-administration (F _(2,1076)_ = 279.4 p < 0.0001 after one way ANOVA). Analysis of the Bonferroni post hoc comparisons showed the decline of analgesic effects of oxycodone after 14 days of extended access of oxycodone self-administration (p < 0.001, Fig. 1D). Finally, oxycodone withdrawal-induced hyperalgesia was assessed using the Von Frey Test. Paw withdrawal responses in response to a mechanical stimulus were measured at the beginning and at the end of the behavioral paradigm (pre- and post-oxycodone, Fig. 1E) and the data were expressed a percentage reduction of force (g) compared to baseline responses. The results showed that HS developed withdrawal-induced hyperalgesia following extended access to oxycodone (t_538_ = 7.41, p < 0.001, Fig. 1E).

### Sex differences on Oxycodone addiction-like behaviors in HS rats

No sex differences were observed during the acquisition phase under ShA conditions. However, during LgA, female HS rats showed higher rates of oxycodone escalation of intake (Fig. 2A). The two-way ANOVA with sex as between factor and sessions as within factor showed a significant effect of sex (F_(1, 542)_ = 6.577; p <0.001), sessions (F_(17, 9210)_ = 224.5; p <0.001) and of the interaction sex*sessions (F_(17, 9210)_ = 5.661; p < 0.001). The Bonferroni post hoc test demonstrated that females self-administered more oxycodone compared to males starting from day 6 of the LgA phase (day 10 on Fig. 2A) until the end of the behavioral protocol (p < 0.01).

**Figure 2.**
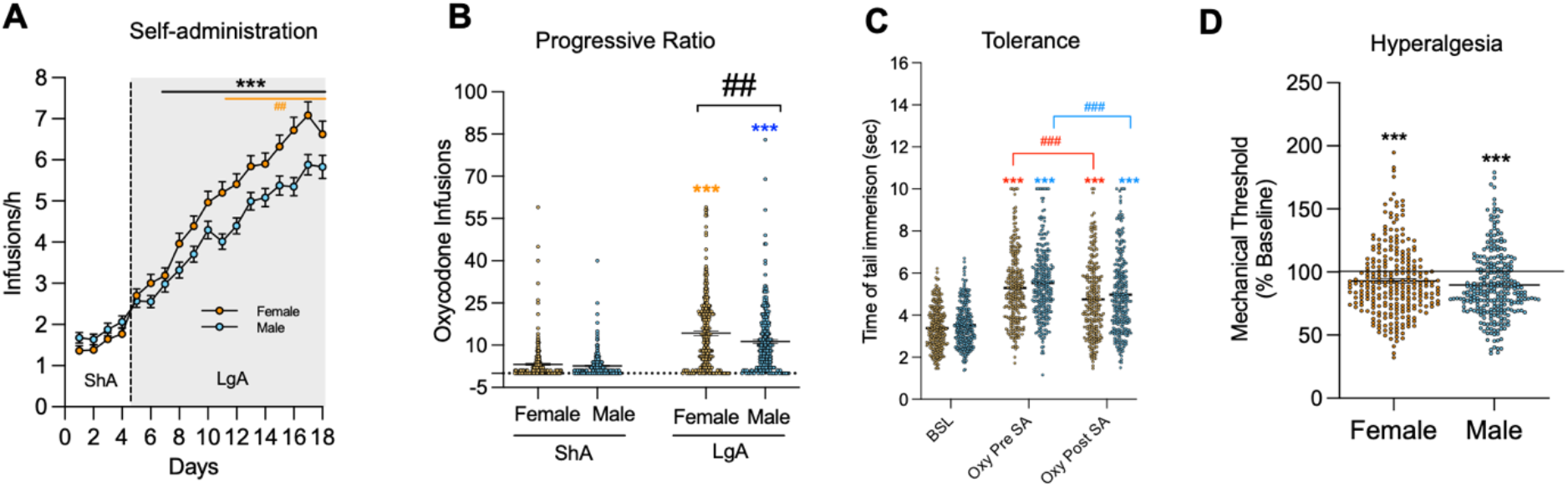
Sex differences in addiction-like behaviors in oxycodone dependent HS rats. A) Oxycodone infusions during short (2h, ShA) and long (6h, LgA) access of oxycodone self-administration, *** p < 0.01 vs LgA 1 and ## p < 0.01 vs males). B) Violin plot of number of oxycodone infusions under progressive ratio (PR) test after ShA and LgA (*** p < 0.01 vs ShA, ## p < 0.01 vs males). C) Development of tolerance to the analgesic effects of oxycodone. Tail withdrawal threshold at baseline, after 3 infusions of oxycodone pre- and post-extended access of oxycodone SA in male and female HS rats (N=523, *** p < 0.001 vs BSL and ### p < 0.001 vs Oxy pre-SA). E) Development of withdrawal induced hyperalgesia in male and female HS rats. Data are expressed as % change from baseline force in the Von Frey test. *** p < 0.001 vs BSL.

In the progressive ratio test, the two-way ANOVA with sex as between factor and time as within factor showed a significant effect of sex (F _(1, 537)_ = 8.28; p < 0.01), time (F_(1, 537)_ = 337.2; p <0.001) and of the interaction sex*time (F_(1, 537)_ = 4.83; p < 0.05). Analysis of the Bonferroni post hoc test confirmed that motivation for oxy was increased in both sexes over time from ShA to LgA (p < 0.001) and that females had significantly higher motivation for oxy compared to males after long access (p < 0.01, Fig. 2B).

Male and female HS Rats showed similar levels of baseline response to the tail immersion test, similar responses to the analgesic effects of oxycodone and similar rates of development of tolerance to the analgesic effects of oxycodone (no treatment * sex interaction, F_(2, 1074)_ = 0.19; p = NS, Fig 2C).

Similarly to tolerance, the development of hyperalgesia was comparable between males and females (no sex differences in paw withdrawal thresholds during withdrawal, (t_537_ = 1.32, p = NS Fig 2D).

### Addiction Index: evaluation of individual differences in addiction-like behaviors

To evaluate addiction-like behaviors in individual rats, we used an Addiction Index (22, 30–33), which includes different addiction-related behaviors: escalation of oxycodone intake under a fixed-ratio (FR) schedule of reinforcement, motivation to maintain responding under a progressive-ratio (PR) schedule of reinforcement, tolerance to the analgesic effects of oxycodone and withdrawal-induced hyperalgesia. To combine these four behavioral outputs, each measure was normalized into an index using its *Z*-score 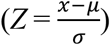 where *χ* is the raw value, *μ* is the mean of the cohort, and *σ* is the standard deviation of the cohort. We thus obtained an FR Index, PR Index, Tolerance Index and Pain Index (Fig. 3A). For the FR Index, the final values were obtained by calculating a Z-score that was the average of the Z-scores of the last 3 days of escalation. For the PR index we calculated the Z-score of the breakpoint, for the tolerance index the Z-score was calculated from the difference in response to the analgesic effects of oxy pre and post extended access of oxy self-administration and for the Pain Index, the Z-score was calculated from the percent reduction of pain thresholds during withdrawal compared with naive baseline. Finally, we calculated the Addiction Index by averaging the Z-scores of the four dependent variables (FR Index, PR Index, Tolerance Index and Pain Index; Fig 3A). We performed a Principal Component Analysis (PCA) on the four addiction-like behaviors (escalation, motivation tolerance and withdrawal-induced hyperalgesia) that compose the Addiction Index to determine if dimensionality could be reduced while maintaining variability. The PCA revealed only one component with an eigenvalue >1, explaining 40% of the variance and to which all four behaviors contributed in a valuable way (*r* = 0.19 to 0.78). This analysis supports the Addiction Index as a reflection of this first principal component and thus as a good approach to capture the variability in addiction-like behavior in one dimension.

**Figure 3.**
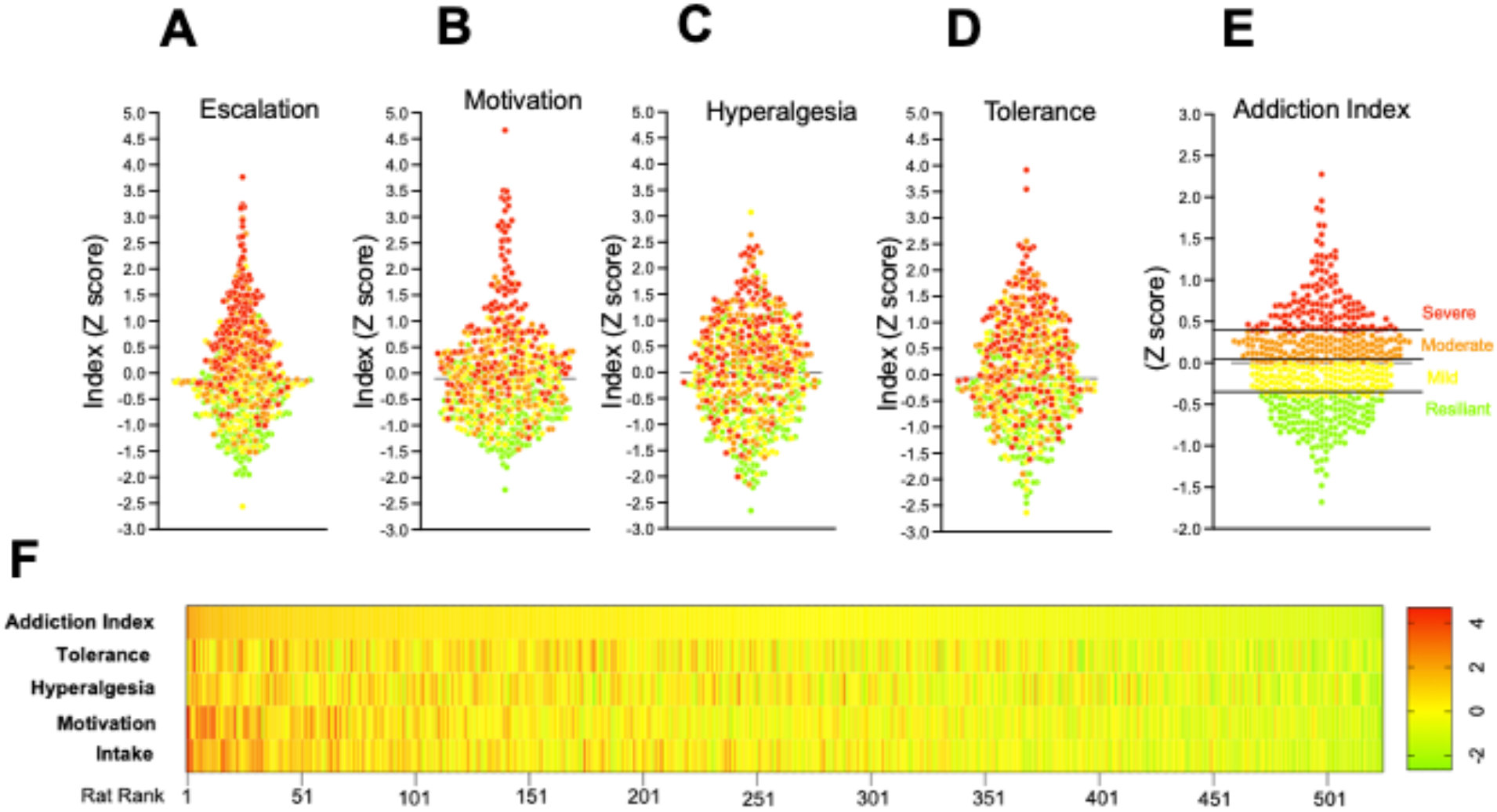
The Addiction Index: a comprehensive evaluation of addiction-like behaviors in HS rats. A) Z-score for escalation, B) motivation, C) hyperalgesia and D) Tolerance in the whole population. E) Addiction Index for animals with severe (red), Moderate (orange), Mild (yellow) and Low (Resilient, green) oxycodone addiction like behaviors. Representation of the individual rats, resilient (green) or vulnerable (red) along the two first principal components, based on escalation, motivation, hyperalgesia and tolerance z-scores. F) Representation of the addiction index for the individual rats with the constituting individual z-scores and their identification as resilient or vulnerable and male or female.

Using the addiction index, animals can be ranked from low to high addiction-like behaviors. We differentiated between animals with low (resilient), mild, moderate and severe addiction like behaviors in the whole population, by dividing them in 4 equal quartiles (Fig 3B).

Oxycodone intake was progressively higher between the resilient, mild, moderate and severe groups. The two-way ANOVA with group as between factor and sessions as within factor showed a significant effect of group (F_(3, 535)_ = 75.85; p < 0.0001), sessions (F_(17, 535)_ = 239.1; p < 0.0001) and of the interaction group*sessions (F_(51, 9095)_ = 18.48; p < 0.0001). Pairwise comparisons for the significant main effects of groups, demonstrated that each of the subgroups (divided based on their Addiction Index) obtained significantly more oxycodone of their immediate lower subgroup (severe > moderate > mild > resilient, Fig 4A). In the progressive ratio test (Fig. 4B), the two-way ANOVA showed significant effects of group (F_(3, 535)_ = 60.04; p < 0.0001), time (F_(1, 535)_ = 404.3; p < 0.0001) and of the interaction group*time (F_(3, 535)_ = 38.7; p < 0.0001). Pairwise comparisons showed that all subgroups (divided based on their Addiction Index), showed increased breakpoint after long access, compared to after short access. In addition, each of the subgroups (divided based on their Addiction Index) showed increased motivation for oxycodone of their immediate lower subgroup (severe > moderate > mild > resilient, Fig 4B)

**Figure 4.**
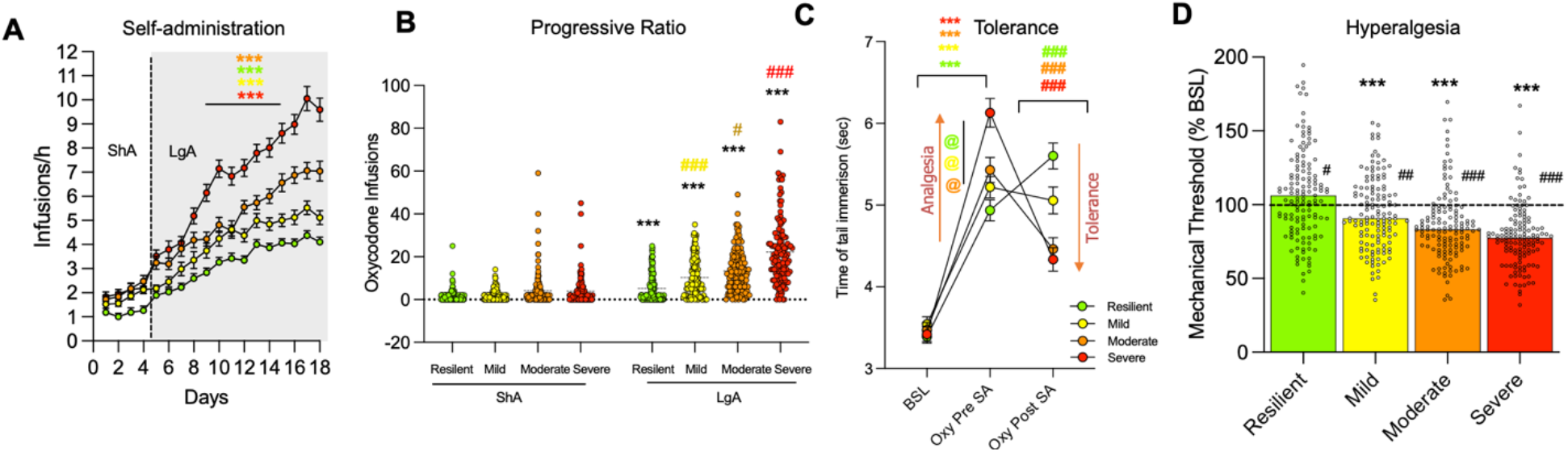
Identification of four degrees of vulnerability to oxycodone addiction-like behaviors in HS rats. A) Oxycodone infusions during short (2h, ShA) and long (12h, LgA) access of oxycodone self-administration (*** p <0.001). B). Number of oxycodone infusions under progressive ratio (PR) test at the end of the ShA and LgA phases (*** p <0.001 vs ShA, # p < 0.05 and ### p < 0.001 vs immediate lower subgroup). C) Development of tolerance to the analgesic effects of oxycodone. Tail withdrawal threshold at baseline, after 3 infusions of oxycodone pre- and post-extended access of oxycodone SA in severe, moderate, mild and resilient HS rats (*** p < 0.001 vs BSL and ### p < 0.001 vs Oxy pre-SA, @p< 0.05 vs immediate lower subgroup). E) Development of withdrawal induced hyperalgesia severe, moderate, mild, and resilient HS rats. Data are expressed as % change from baseline force in the Von Frey test. *** p < 0.001 vs BSL. # p < 0.05, ## p < 0.01 and ### p < 0.001 vs immediate lower subgroup.

In the tail immersion test, the two-way ANOVA showed a significant effect of the group*time interaction (F_(6, 1070)_ = 22.00; p < 0.0001). The Bonferroni post doc test demonstrated that all the groups showed increased analgesia after the first injection of oxycodone (p<0.001, Fig 4C). Interestingly, the severe group showed higher rates of analgesia compared to all the other groups (p<0.001, Fig 4C). After extended access of oxycodone self-administration, moderate and severe animals showed development of tolerance to the analgesic effects of oxycodone (as demonstrated by the decreased response in the tail immersion test after a dose of oxycodone compared to the response pre-self-administration). On the other hand, the resilient group showed a significantly increased response compared to the baseline pre-self-administration, suggestive of a sensitization of the analgesic effect of oxycodone that was probably due to the low and intermittent level of oxycodone intake during extended access to oxycodone self-administration (Fig 4C). This effect in the resilient group extended also to the Von Frey test. The analysis showed that all the groups showed emergence of hyperalgesia during acute withdrawal from oxycodone (p < 0.001 vs BSL) except for the resilient group that showed instead a significantly increased paw withdrawal response in the Von Frey test after extended access of oxycodone self-administration (p < 0.05 vs BSL). The 1 way ANOVA showed a significant difference between the groups (F_(3, 535)_ = 30.64; p < 0.0001). The Bonferroni post doc test demonstrated that each of the subgroups (divided based on their Addiction Index) showed increased oxycodone withdrawal-induced hyperalgesia compared to their immediate lower subgroup (severe > moderate > mild > resilient, Fig 4D)

## Discussion

We characterized addiction-like behaviors in over 530 genetically diverse HS rats by establishing a large behavioral screening program to study different oxycodone-related behaviors (escalation, motivation, tolerance, and withdrawal-induced hyperalgesia). We used extended access to intravenous self-administration combined with behavioral characterization of motivation using progressive ratio responding, development of tolerance to the analgesic effects of oxycodone (tail immersion test), and development of withdrawal-induced hyperalgesia (von Frey test). The extended access to the self-administration model with the escalation of drug use is highly relevant to opioid use disorders (34, 35) and is associated with brain neuroadaptations that have been identified in humans with drug use disorders (36–40). Compelling evidence indicates that the escalation of oxycodone intake using this protocol is associated with compulsive oxycodone use, measured by the escalation of intake (17) and an increase in PR responding (17). Several studies demonstrated that the escalation model has been shown to exhibit 7 of the 11 items in the DSM-5, including most of the criteria that are required for severe use disorder: tolerance (41), withdrawal (42, 43), escalation of intake (44), inability to quit (45, 46), (5) substantial time spent to obtain the drug (47), (6) important social work or recreational activities abandoned because of drug use (37, 48), and (7) continued use despite adverse consequences (40, 49–51). Previous studies have employed smaller numbers of animals, used inbred strains that were selectively bred to be high or low responders (52) and used the BXD recombinant inbred mouse panel (53) or Hybrid Mouse Diversity Panel (HMDP) (54). Moreover, several of these studies also used other proxies for addiction-like behaviors, like exploratory locomotion (52) or drug-induced motor effects (55) following passive drug injections.

This behavioral screening is the first high-throughput intravenous oxycodone self-administration study in HS rats, a genetically diverse population that self-administers oxycodone with high inter-individual variability and low intra-individual variability. Such behavioral variability between subjects and consistency within subjects will maximize the identification of gene variants associated with compulsive oxycodone use and oxycodone dependence.

The evaluation of different addiction-like behaviors is essential as multiple elements of addiction vulnerability were found to be independently heritable (56), and are likely controlled by distinct genes that remain unidentified. Opioid self-administration and the psychoactive effects of opioids in rodents, like opioid use disorder in humans, have a heritable component. Inbred strain studies have identified strain differences in baseline heroin self-administration (57-59), escalation of heroin self-administration (60), preferred dose of opioids (60), acute reactivity to opioid withdrawal (61, 62), antinociceptive effect of opioids (63), opioid-induced conditioned place preference(64), brain levels of proenkephalin (57), and μ opioid receptor regulation (65). Thus, there is ample evidence of heritable differences in response to opioids in various inbred rat strains. In our work, we found that escalation, motivation, tolerance, and withdrawal-induced-hyperalgesia were typically very similar within-subject, while there was large variability between subjects (Fig. 4). By computing an Addiction Index, we could better characterize the addiction-like behaviors in the whole population by distinguishing their addiction phenotype into resilient, mild, moderate, and severe addiction-like behaviors. Resilient animals showed a low escalation of intake and lower motivation and did not develop tolerance to the analgesic effects of opioids and withdrawal-induced hyperalgesia. Severe animals showed an extreme escalation of intake, high motivation, hyperalgesia, and tolerance. Interestingly, the severe group seemed to have a higher sensitivity to the analgesic effects of oxycodone, with very high levels of oxycodone analgesia in the tail immersion test before extended access to oxycodone self-administration. However, the severe group also showed the highest levels of tolerance to the oxycodone analgesic effects, which could be explained by the very high levels of drug intake during the extended access to the oxycodone self-administration paradigm.

Future studies will be designed to elucidate the correlation between the first analgesic response to oxycodone and the risk of developing addiction-like behaviors.

Data from the human literature suggest that despite minor differences in the level of hepatic enzymes involved in the metabolism of oxycodone between males and females, there is no significant difference in the pharmacokinetics or pharmacodynamics of oxycodone use (66). However, converging evidence suggests that females show increased adverse events like nausea or vomiting compared to men (66). In human studies, it has been reported that the abuse of, and craving for, prescription pain medications and opioids are more frequent in women (67). The results presented here and our recent work (23) confirmed these findings by showing that female rats self-administered more oxycodone than male rats. Interestingly, in a short-access model of oxycodone self-administration, female rats also self-administered more oxycodone at higher doses than males. However, no sex differences in brain or plasma levels of oxycodone were found with a low-dose infusion of oxycodone (25). Similar to (25) who used a low-dose infusion of oxycodone, our previous work found no sex differences in plasma oxycodone levels after a high-dose infusion of oxycodone (23), but a significant elevation of brain oxycodone levels in male rats compared with female rats at 30 min (23). These data suggest that the higher level of oxycodone self-administration that was observed in females may reflect behavioral compensation for lower brain oxycodone levels in females, which could be attributable to sex-related differences in the distribution characteristics of oxycodone. Our findings are also consistent with human pharmacokinetic data showing lower oxycodone levels in females (68–70). These results demonstrate that our animal model of oxycodone escalation is highly relevant to the human condition and highlights sex differences in oxycodone-related behaviors.

The behavioral characterization of oxycodone addiction-like behaviors in HS rats was the first step in characterizing and understanding individual differences in a genetically diverse population. Future studies will combine these phenotypic data with genetic and genomic data obtained from these animals through whole genome sequencing to perform a GWAS to identify gene variants that contribute to vulnerability and resilience to oxycodone addiction-like behaviors. This GWAS study will then allow for the identification of specific genetic loci, estimation of heritability, investigation of genetic correlations, performance of phenome-wide association studies (PheWAS) and transcriptome-wide association studies (TWAS). The results from these studies might provide valuable information on the genetic behind individual differences in the vulnerability and resilience to oxycodone abuse and might also provide several addiction-like traits to be used for personalized diagnoses and/or novel treatment approaches. Finally, to facilitate investigating the biological origins of the difference in oxycodone addiction-like behaviors through epigenetic, transcriptomic (55), microbiome, neurobiological, or other mechanisms in HS rats beyond genetic factors, samples of these behavioral and genetically characterized animals were collected and made available free of charge through the oxycodone biobank (32).

## Acknowledgements

This work was supported by National Institutes of Health grant DA044451 from the National Institute on Drug Abuse. The authors declare no competing financial interests.

## Notes

Funding and disclosure, The authors declare no conflict of interest.

### Competing Interest Statement

The authors have declared no competing interest.

